# A Single Dose Of Ayahuasca Modulates Salivary Cortisol In Treatment-Resistant Depression

**DOI:** 10.1101/257238

**Authors:** Ana Cecília de Menezes Galvão, Raíssa Nóbrega de Almeida, Erick Allan dos Santos Silva, Fúlvio Aurélio de Morais Freire, Fernanda Palhano-Fontes, Heloisa Onias, Emerson Arcoverdee, João Paulo Maia-de-Oliveira, Draúlio B Araújo, Bruno Lobão-Soares, Nicole Leite Galvão-Coelho

**Affiliations:** Laboratory of Hormone Measurement, Departament of Physiology, Federal University of Rio Grande do Norte, Natal, RN, Brazil; Postgraduate Program in Psychobiology, Federal University of Rio Grande do Norte, Natal, RN, Brazil; National Institute of Science and Technology in Translational Medicine, Natal, RN, Brazil; Brain Institute, Federal University of Rio Grande do Norte, Natal, Brazil; Onofre Lopes University Hospital, Federal University of Rio Grande do Norte, Natal, Brazil; Departament of Biophysics and Pharmacology, Federal University of Rio Grande do Norte, Natal, Brazil; Departament of Clinical Medicine, Federal University of Rio Grande do Norte, Natal, RN, Brazil

**Keywords:** ayahuasca, awakening salivary cortisol response, plasma cortisol, treatment-resistant depression, hypocortisolemia

## Abstract

Major depression is a highly prevalent mood disorder, affecting about 350 million people, and around 30% of the patients are resistant to currently available antidepressant medications. Recent evidence from a randomized placebo-controlled trial supports the rapid antidepressant effects of the psychedelic ayahuasca in treatment-resistant depression. The aim of this study was to explore the effect of ayahuasca on plasma cortisol and awakening salivary cortisol response, in the same group of treatment-resistant patients and in healthy volunteers. Subjects received a single dose of ayahuasca or placebo, and both plasma and awakening salivary cortisol response were measured at baseline (before dosing) and 48h after the dosing session. Baseline assessment (D0) showed blunted awakening salivary cortisol response and hypocortisolemia in patients (DM), both with respect to healthy controls group (C). Salivary cortisol also was measured during dosing session and we observed a large increased for both C and DM that ingested ayahuasca, than placebo groups. After 48h of the dosing session (D2) with ayahuasca, awakening salivary cortisol response (for both sexes) of treated patients became similar to levels detected in controls. This was not observed in patients that ingested placebo. No changes in plasma cortisol were observed after 48 hours of ayahuasca or placebo ingestion for both groups and sexes. Therefore, these findings point to new evidence of modulation of ayahuasca on salivary cortisol levels, as cortisol acts in regulation of distinct physiological pathways, emotional and cognitive processes related to etiology of depression, this modulation could be an important part of the antidepressant effects observed with ayahuasca. Moreover, this study highlights the importance of psychedelics in the treatment of human mental disorders.

## 1 INTRODUCTION

Major depression is a highly prevalent mood disorder, affecting about 350 million people worldwide (1). It is more prevalent in women than men and has a huge impact on general health of the patients (2, 1).

Major depressive disorder (MDD) has been closely associated with deregulations of the hypothalamic-pituitary-adrenal (HPA) axis, both at rest and in response to stress (3, 4, 5). Some studies report changes in cortisol response that occur soon after awakening (6). Most often reported, cortisol awakening response (CAR) is increased in patients with major depression suggesting hyperactivity of the HPA axis (7). However, there is increasing evidence of hypocortisolism in patients with depression, which has been interpreted as an indication of HPA axis fatigueness in response to recurrent depressive episodes (8, 9). Such discrepancies can be attributed to a number of factors including subtypes of depression, depression severity, sex, duration of illness and socioeconomic status (10, 11, 12, 13).

Cortisol assessments have also served as an important biomarker of treatment response. For instance, patients with depression who responded to an 8-week treatment with fluoxetine, a selective serotonin reuptake inhibitor, presented decreased levels of cortisol (14). At present, the large majority of currently available antidepressants take about two weeks for the beginning of their therapeutic effects (15, 16, 17, 18).

Recently, however, psychedelics have been emerging as a promising fast-acting antidepressant (19, 20, 21, 22, 23). Clinical trials have pointed to a positive effect of psychedelics in depression. A recent open label trial in treatment-resistant depression observed a reduction of up to 87% in depression severity, already at 24h after a single dosing session with ayahuasca (24, 23). Ayahuasca was originally used for medicinal purposes by indigenous populations groups in Brazil, Ecuador, Peru and Colombia, and later its ritualistic use became more popular by its presence in ceremonies of different syncretic churches in Brazil, which is currently spreading to other parts of the world (25).

Ayahuasca is a decoction of a mixture of two plants: *Psychotria viridis* and *Banisteriopses caapi* (26). *P. viridis* contains the psychedelic tryptamine N,N-dimethyltryptamine (N,N-DMT), whose action is mediated by serotonin (5-HT2A) and sigma-1 receptors (27, 28, 29, 30). *B. caapi* contains β-carbolinic alkaloids (harmaline, harmine and tetrahydroharmine), which work as indirect monoaminergic agonist due to the inhibition of monoamine oxidase isoenzyme (MAO) (31, 32, 33). Regular users of ayahuasca in religious contexts have shown low level of psychopathologies (34, 35), low scores on the state scales related to panic and hopelessness (36) as well as good performances in cognitive neuropsychological tests (37, 38). Moreover, this brew does not exhibit dose tolerance, i.e., the decrease of the effect of a drug or medication by excessive or frequent exposure of the patient to its active principle, and is not addictive (39, 40, 41).

Considering that the main neurobiological actions of ayahuasca are strongly related to key physiological systems altered in major depression, and taking into account the low incidence of mental disorders in regular users in religious context, and previous results from open label trial (23), we recently conducted a randomized placebo-controlled trial with ayahuasca in patients with treatment-resistant depression. Our results suggest significant and rapid reduction in depressive symptom one day after a single ayahuasca dose, when compared to placebo (42). Herein, we explored the effects of ayahuasca on the salivary cortisol awakening response and plasma cortisol, in patients with treatment-resistant depression and in healthy individuals.

Our hypotheses are that patients and controls will show, in baseline, different levels of plasma cortisol and awakening salivary cortisol response and the cortisol levels in patients will be correlated with severity and/or duration of disease. Moreover, ayahuasca, but not placebo, will increase cortisol levels acutely (43), during dosing session and after 48 hours of its ingestion in volunteers patients and control, but with different intensity. The responses will be correlated with improvement in depression symptoms in patients group (42).

## 2 METHODS

This is a randomized double-blinded placebo-controlled trial using a parallel arm design. Patients were referred from psychiatric units of the Onofre Lopes University Hospital, in Natal/RN, Brazil, and through media and internet advertisements. All procedures took place at the University Hospital. The study was approved by the Research Ethics Committee of the University Hospital (# 579.479; see supplementary material), and all subjects provided written informed consent prior to participation. This study is registered in http://clinicaltrials.gov (NCT02914769).

### 2.1 Volunteers

Seventy-one volunteers participated in the study: 43 healthy volunteers, control group (C), (19 men and 24 women) without history or diagnosis of major illness or psychiatric disorders, and 28 patients, major depression (DM), (7 men and 21 women) with treatment-resistant depression, defined as those with inadequate responses to at least two antidepressants from different classes (44). Patients were screened for exclusion due to previous experience with ayahuasca, current medical disease based on history, pregnancy, current or previous history of neurological disorders, history of schizophrenia or bipolar affective disorder, history of mania or hypomania, use of substances of abuse, and suicidal risk. Selected patients were in a current moderate to severe depressive episode at screening by the Hamilton Depression Rating Scale (HAM-D≥17). Depressive symptoms were monitored at baseline and two days after the dosing session by a clinical scale traditionally used to measure depression severity: the Montgomery-Åsberg Depression Rating Scale (MADRS). All patients were not using any antidepressant medication during the trial, however they all were under regular use of benzodiazepines.

Volunteers from both groups (healthy and patients) were randomly assigned (1:1) to receive ayahuasca or placebo using 10-gauge blocks. Half of the patients and half of the controls received ayahuasca while the other half received placebo. All investigators and patients were blinded to the intervention assignment.

### 2.2 Ayahuasca and placebo

The substance used as placebo did not have psychoactive properties, but induced a light gastrointestinal discomfort, and simulated some organoleptic properties of ayahuasca. It is a brown liquid with a bitter and sour taste. It contained water, yeast, citric acid, zinc sulfate and a caramel dye.

A single batch of ayahuasca was used throughout the study. It was prepared and supplied free of charge by a branch of the Barquinha church, based in the city of Ji-Paraná, Brazil. The alkaloid concentrations in the ayahuasca batch were analyzed by mass spectroscopy twice during the trial. On average, the ayahuasca contained (mean±DP): 0.36±0.01 mg/mL of N,N-DMT, 1.86±0.11 mg/mL of harmine, 0.24±0.03 mg/mL of harmaline and 1.20±0.05 mg/mL of tetrahydroharmine (THH).

### 2.3 Salivary cortisol

Saliva was collected using a specific cotton stick called Salivette (Sarstedt, Germany). Volunteers were instructed to place the cotton in the mouth without touching and masticating it for a period of about 1 to 2 minutes. Before and during collection subjects remained at rest and no liquid or food were allowed.

Saliva samples were stored at −80°C in the Laboratory of Hormonal Measures (UFRN) and the salivary cortisol was measured using the ELISA DGR – SLV 4635 kit (DGR International, Inc, Germany).

### 2.4 Plasma cortisol

Blood samples were collected in the morning (7:00 a.m), for total plasma cortisol (PC) assessment. All volunteers were at fast and at complete rest for 45 minutes prior to the exam. After, the samples were stored at −80°C in the Laboratory of Hormonal Measures (UFRN). Total plasma cortisol was measured by ELISA using the DGR-SLV 1887 kit (DGR International, Inc, Germany).

### 2.5 Experimental Procedure

Figure 1 shows the experimental design of the study. After admission, volunteers were interned in the psychiatry division of the University Hospital (HUOL) one night before dosing (D-1), when the MADRS scale of depression was applied. The volunteers slept in the hospital. At 6:00 a.m next day (D0) saliva samples were collected for measuring awakening salivary cortisol and at 7:00 a.m the blood samples were collected for PC assessment. The procedure of collecting of awakening salivary cortisol response consisted of 3 saliva samples: (i) at awakening (+5 minutes), (ii) +30 minutes, and (iii) +45 minutes later.

**Figure 1.**
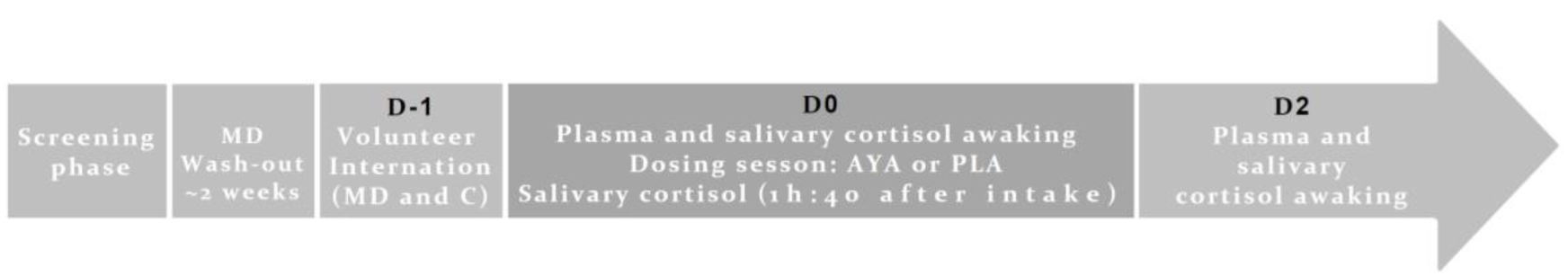
Temporal line of experimental design. In this study, patients with major depression (MD) were selected by clinicians in the screening phase that was followed by a pharmacological wash-out phase of approximately two weeks. In D-1, all subjects, healthy volunteers of control group (C) and MD, slept at the hospital (internation) and plasma and salivary cortisol at waking time, at D0, were collected. Also in D0 the dosing session happened, and subjects (C and MD) were divided in two subgroups each, depending on placebo (PLA) or ayahuasca (AYA) intake. During dosing session saliva was collected for cortisol measurement, 1h40 after intake. On D2 (48 hours after intake), again, plasma and saliva collection at awake were performed for cortisol dosage.

After a light breakfast, volunteers received instructions and guidance on the effects they could experience after taking ayahuasca, and strategies to help alleviating any difficulties encountered.

Dosing session started around 10:00 a.m. They received a single dose of 1 ml/kg of ayahuasca (AYA) adjusted to contain 0.36 mg/kg of N,N-DMT, or 1 ml/kg of placebo (PLA). During the entire session subjects were asked to remain quiet with their eyes closed while concentrating on their body, thoughts and emotions. They were allowed to listen to a pre-defined music playlist. Volunteers were supported by at least two researchers offering assistance when needed. Acute response (%) of salivary cortisol were assessed during dosing session at two instants: (i) immediately before dosing, and (ii) +1h40 minutes after the ingestion of placebo or ayahuasca.

On the following day (D1), volunteers slept again in the hospital, and when woken up at 6:00 a.m. of the next day (D2), 48h after dosing session, 3 saliva samples were collected for measuring awakening salivary cortisol and at 7:00 a.m the blood samples were collected for PC assessment. Again, the MADRS scale of depression was applied.

### 2.6 Statistical analysis

Statistical analysis was conducted in Statistic 12.5 (data analysis software system), and the level of significance was set at p < 0.05 for all tests. Graphics were built in R 3.4.1 (RStudio).

The area under the curve (AUC) was calculated from the 3 points of salivary cortisol at waking time. Both salivary and plasma cortisol levels were normalized by the logarithm to use parametric tests.

A parametric test of Analysis of Covariance (ANCOVA) was used to analyze differences between groups (healthy and patients) at baseline, for both salivary (AUC of awakening salivary cortisol) and plasma cortisol. Sex was inserted as co-variable.

At baseline, *Spearman* correlations were calculated across plasma cortisol and AUC of awakening salivary cortisol of patients and controls and scores of scales of depression (HAM-D and MADRS) and duration of disease of patients.

General Linear Models (GLM) and Fisher *post-hoc* tests were used to evaluate interaction among: changes of AUC of awakening salivary cortisol response along the days (D0 and D2), which was considered as dependent variable, and sex (men and women), groups (healthy volunteers and patients) and treatment (AYA or PLA) as independent variable. For plasma cortisol, this same analysis was applied but sex was not used as independent variable, because the number of male patients who received placebo was too small (n=2).

Acute response (%) of salivary cortisol during the dosing session were evaluated 1h40 after ayahuasca or placebo ingestion and assessed by Mann-Whitney test.

Moreover, *Spearman* correlations test were calculated across acute response (%) of salivary cortisol during the dosing session for controls and patients of AYA and PLA groups, plasma cortisol and AUC of awakening salivary cortisol of D2 for patients and controls of each treatment and scores of MADRS for patients of each treatment.

## 3 RESULTS

Socio-demographic characteristics of healthy volunteers, control group (C), and patients with major depression (MD) are summarized in table 1. All volunteers (n=71; MD=28, C=43) were Brazilian, adults (MD= 41.54 ± 11.55, C= 31.21 ± 9.87 years, t (69) =4.03 p<0.0001). Patients showed significant lower socioeconomic status backgrounds than healthy volunteers: large part of MD was unemployed (MD=54%, C=12%, X^2^(1)= 4.74, p<0.0001), living in a low-income household earning, earn up to 5 minimum wages (MD=87%, C=71%, X^2^(3)=14.03, p=0.003) and had low education, with up to 8 years formal education (MD=39%, C=7%, X^2^(3)=19.88, p=0.0002).

**Table 1.** Socio-demographic characteristics of Seventy-one volunteers participated in the study: 43 healthy volunteers, control group (19 men and 24 women) without history or diagnosis of major illness or psychiatric disorders, and 28 patients with major depression (7 men and 21 women).

|  | Controls | Patients | Statistical analysis |
| --- | --- | --- | --- |
| Participants, n | 43 | 28 |  |
| Age (years) | 31.21 ± 9.87 | 41.54 ± 11.55 | t(69)=4.03 p < 0.0001 |
| Gender (M/F) | 19/24 | 7/21 | X<sup>2</sup>(1) = 2.69 p =0.10 |
| Unemployed (%) | 5/43 (12) | 15/28 (54) | X<sup>2</sup>(1) = 14.74 p < 0.0001 |
| Household income |  |  | X<sup>2</sup>(3) = 14.03 p = 0.003 |
| < 5 wages (%) | 19/86 (71) | 22/56 (87) |  |
| 6-10 wages (%) | 14/43(7) | 2/28 (6.6) |  |
| 11 or more wages (%) | 10/43 (21) | 4/28 (6.6) |  |
| Education |  |  | X<sup>2</sup>(3) = 19.88 p = 0.0002 |
| Up to 8 years, n (%) | 3/43 (7) | 11/28 (39) |  |
| 9-11 years, n (%) | 4/43 (9) | 8/28 (29) |  |
| 12-16 years, n (%) | 18/43(42) | 4/28 (14) |  |
| 17 or more years, n (%) | 18/43 (42) | 5/28 (18) |  |

On average, patients presented 11.03±9.70 years of depressive symptoms and met criteria for moderate-to-severe depression (HAM-D =21.83±5.35). Usually, they were treated previously with 3.86±1.66 different types of antidepressants and two patients used electroconvulsive therapy as treatment. Majority of patients presented comorbidity, such as personality disorder (76%) and anxiety disorder (31%).

### 3.1 Baseline assessments (D0)

Figure 2 shows the cortisol levels at baseline (D0). Figure 2a shows the AUC of awakening salivary cortisol response for both groups (C and MD) at baseline. AUC level at baseline was lower for patients (MD; n= 20 μ_AUC_=49.4±8.3 cm^2^) than healthy controls (C; n= 41 μ_AUC_=62.5±6.3 cm^2^), and these differences were independent of sex (ANCOVA main effects: Group*: F=9.75 df=1 p=0.002, Sex*: F=0.42 df=1 p=0.51). Figure 2b shows the results for plasma cortisol. The same profile is observed in plasma cortisol at baseline, which was lower in patients (MD; n= 28 μ_PC_=15.12±1.73 mg/dl) than in healthy controls (C; n= 43 μ_PC_=19.52±1.37 mg/dl). Again, these differences were independent of sex (ANCOVA main effects: Group*: F=4.71 df=1 p=0.03, Sex*: F=0.89 df=1 p=0.34). Figure 2b also illustrates patients that show relative (n=17 and 61%) and true hypocortisolemia n=6 and 22.22%), with total plasma cortisol levels below 15μg/dl and 10μg/dl, respectively.

**Figure 2.**
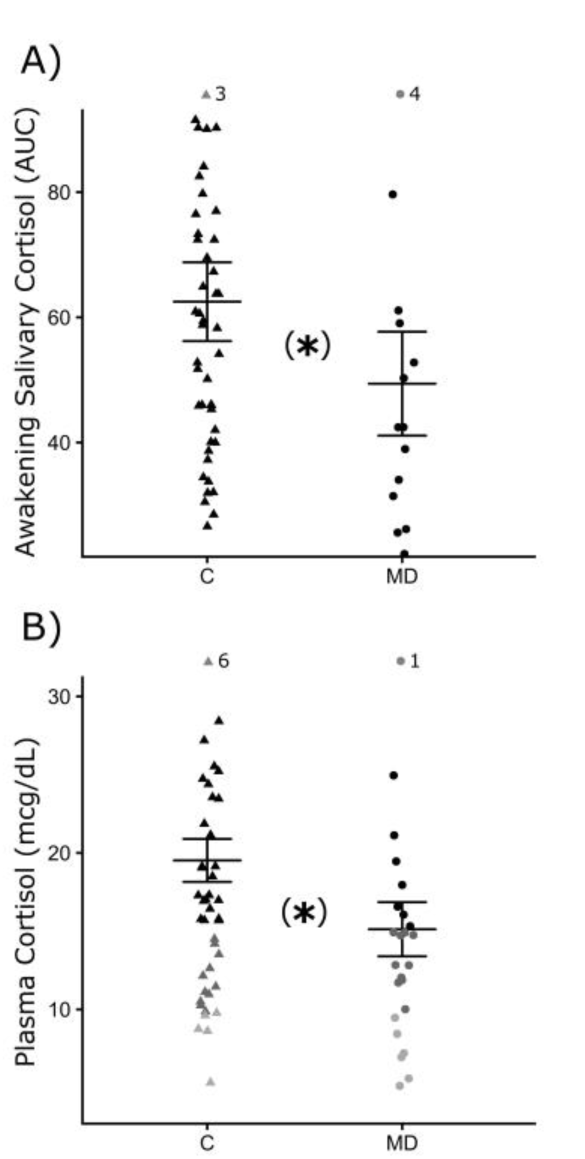
Mean and standard deviation of cortisol levels at baseline (D0) for: A) AUC (area under the curve) of awakening salivary cortisol for control group (C – closed triangle) and patients with major depression (MD – closed circle) and B) plasma cortisol for C and MD. Relative hypocortisolemia (< 15mcg/dl) = dark gray symbols and true hypocortisolemia (< 10mcg/dl) = light gray symbols. Each symbol (triangle or circle) indicate individual value of volunteer. * = statistically significant difference between the groups. ANCOVA, p ≤0.05.

In baseline, the scores MADRS of patients were 32.67±6.31. A positive significant correlation was observed between cortisol levels (plasma and AUC) for controls (p<0.05 *r*_*s*_=0.54), but not for patients. No significant correlations were found across cortisol levels (plasma and salivary), scores of scales of depression (HAM-D and MADRS) and duration of disease for patients (see table 1 of suppl. material for details).

### 3.2 Acute effects of ayahuasca during dossing session

Figure 3 shows the acute response (%) of salivary cortisol observed 1h40 after ayahuasca or placebo ingestion, during dosing session. Figure 3a shows that patients in the ayahuasca group (n=10) presented greater salivary cortisol increases (median=98.72; Q25%=37.89; Q75%=177.16) compared to the placebo group (n=12) (median=23.26; Q25%=-5.44; Q75%=41.65) (Mann-Whitney test U=27 p=0.03). Figure 3b shows the same profile for the healthy volunteers. Controls of the ayahuasca group (n=21) showed greater increases of salivary cortisol levels (median=146.87; Q25%=53.32; Q75%=211.84) compared to the placebo group (n=20) (median=38.50; Q25%=10.33; Q75%=60.27) (Mann-Whitney test U=84 p=0.01). Figure 3c compares patients and controls that ingested ayahuasca, both showing similar changes of salivary cortisol at 1h40min after ingestion of ayahuasca (Mann-Whitney test U=85 p=0.66; patients median=98.72; Q25%=37.89; Q75%=177.16, controls median=146.87; Q25%=53.32; Q75%=211.84). Changes of salivary cortisol (%) for patients from ayahuasca or placebo group during dosing session were not correlated with scores of MADRS at D2 (see table 2 of suppl. material for details).

**Figure 3.**
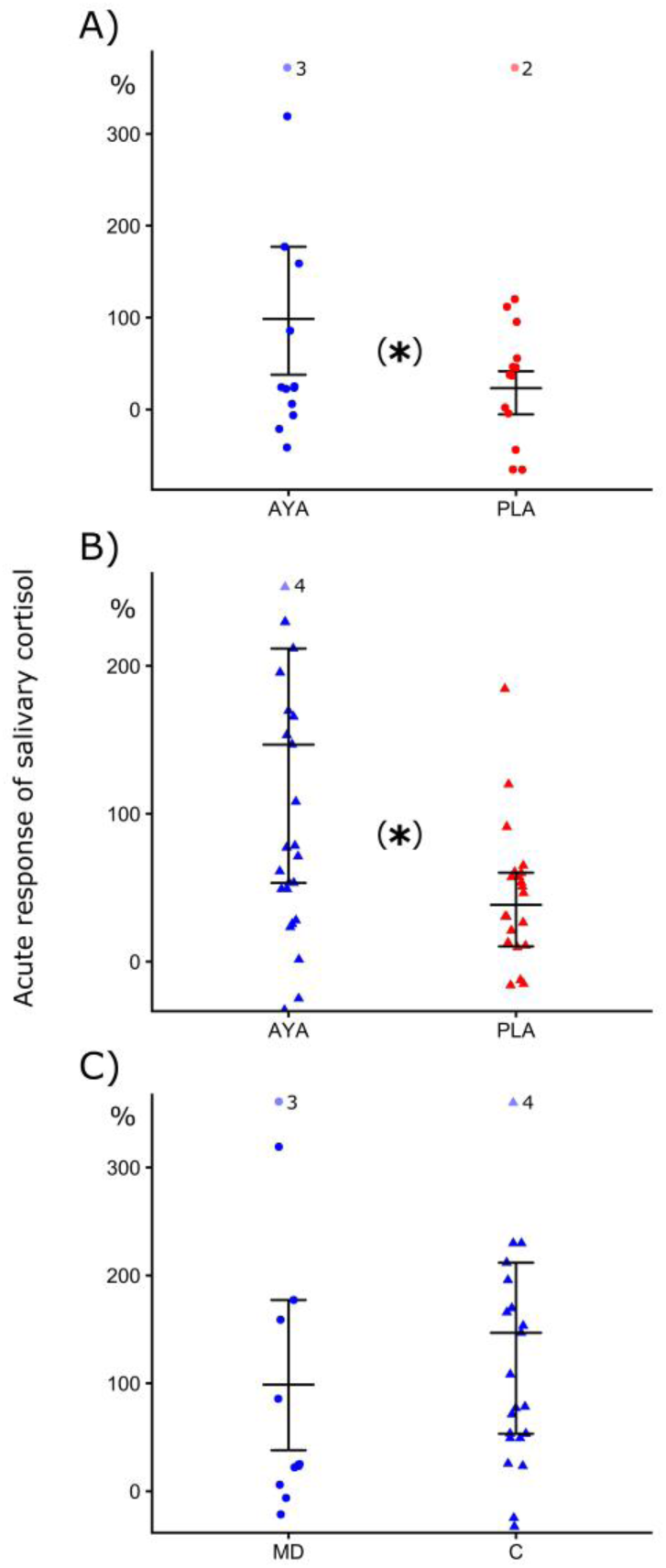
Mean and standard deviation of acute response (%) of salivary cortisol at 1h40min after the dosing session. A) For patients with major depression (MD – closed circle) after ingestion of ayahuasca (AYA-blue color) or placebo (PLA-red color), B) control group (C-closed triangle) after ingestion of of ayahuasca or placebo and C) MD and C after ingestion of ayahuasca. Each symbol (triangle or circle) indicate individual value of volunteer.* = statistically significant difference between the groups. Mann-Whitney non-parametric test, p ≤0.05.

### 3.3 Post-treatment assessments (D2)

Two days after dosing (D2), the scores of MADRS were 55.79±32.14 for patients of ayahuasca group and 61.42±25.7 for patients of placebo group, 77% of patients responded in the ayahuasca group and 64% in the placebo.

Figure 4 shows AUC of awakening salivary cortisol response for both groups (C and MD) and treatments (ayahuasca and placebo) in baseline (D0) and 48h after dosing session (D2) (GLM: Group*Treatment*Days: F=4.57, p=0.03, all values of main effects and interactions of GLM are in table 3 of supplementary material). Figure 4a comparing changes in AUC along the days (D0 and D2) within groups and treatment and no significant variations were found (see table 4 of suppl. material for details of statistical values of Fisher *post-hoc* test). Figure 4b comparing AUC levels across group of patients and control for both treatments, at D2. Was found similar AUC in patients who ingested ayahuasca (μ_AUC_=59.4±12.7 cm^2^) and healthy subjects that ingested ayahuasca (μ_AUC_=50.1±8.2 cm^2^) (Fisher *post-hoc:* p=0.45) or placebo (μ_AUC_=61.9±8.9 cm^2^) (Fisher *post-hoc:* p=0.14). On the other hand, patients that ingested placebo continued presenting lower AUC (μ_AUC_=41.2±14.8 cm^2^) relative to controls that ingested placebo (μ_AUC_=61.9±8.9 cm^2^) (Fisher *post-hoc:* p=0.03), as was observed in baseline. Influence of sex was not observed in statistical analysis (see table 3 of supplementary material). Individual changes of AUC between D0 and D2 were illustrated for each group and treatment in supplementary material (figure 1).

**Figure 4.**
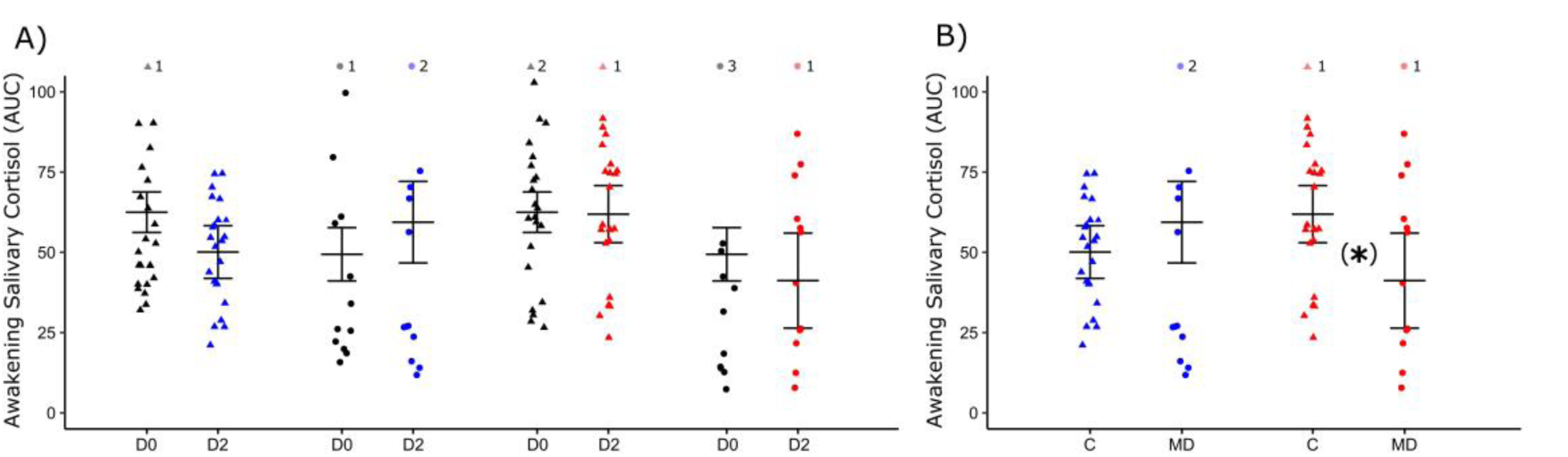
Mean and standard deviation of area under the curve (AUC) of awakening salivary cortisol. A) In baseline (D0-black color) and 48h after dosing session (D2) of control group (C closed triangle) and patients with major depression (MD - closed circle) that ingested ayahuasca (blue color) or placebo (red color). B) AUC in D2 for patients and control and patients of both treatment. Each symbol (triangle or circle) indicate individual value of volunteer. * = statistically significant difference between groups. GLM and post hoc Fisher, p <0.05.

Figure 5 shows the levels of total plasma cortisol for both groups (C and MD) and treatments (ayahuasca and placebo) in baseline (D0) and 48h after dosing session (D2), no significant changes between D0 and D2 within both groups and treatment were observed. Moreover, no significant difference between groups and treatments were found (all values of main effects and interactions of GLM are in table 5 of supplementary material). Individual changes in PC between D0 and D2 were illustrated for each group and treatment in figure 2 of supplementary material.

**Figure 5.**
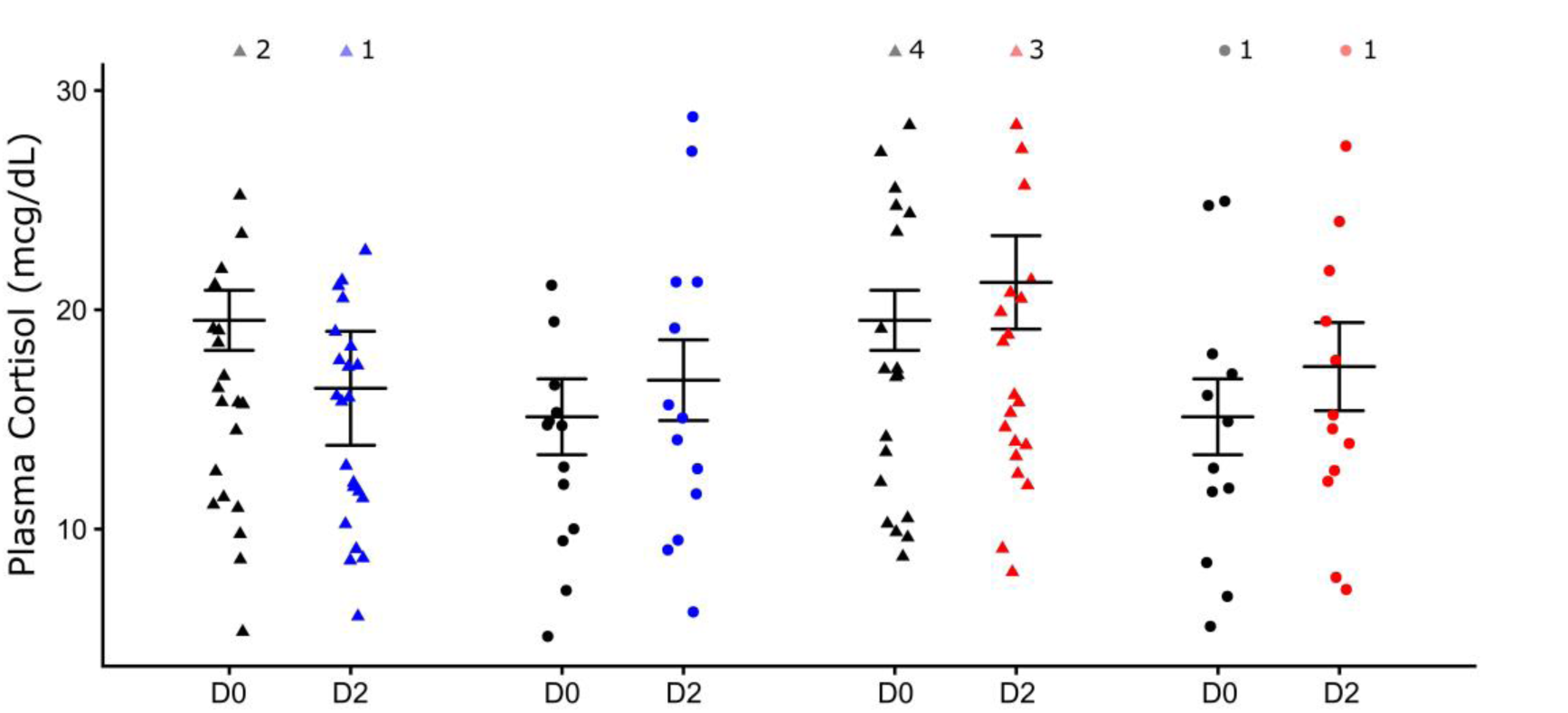
Mean and standard deviation of plasma cortisol (PC) in baseline (D0-black color) and 48h after dosing session (D2) of control group (C – closed triangle) and patients with major depression (MD – closed circle) that ingested ayahuasca (blue color) or placebo (red color). Each symbol (triangle or circle) indicate individual value of volunteer. GLM and *post hoc* Fisher, p ≤0.05.

No statically significant correlations were observed across plasma cortisol and AUC of awakening salivary cortisol of D2 for patients and controls of each treatment and scores of MADRS for patients of each treatment. All values of *Spearman* correlation are in supplementary material, table 2.

## 4 DISCUSSION

In this study we found basal hypocortisolemia and blunted awakening salivary cortisol response in treatment-resistant patients with major depression, compared to healthy subjects. After treatment of patients and controls with ayahuasca or placebo, was observed (1 hour and 40 minutes after ingestion) major acute increases in salivary cortisol of groups that ingested ayahuasca compared to placebo-ingesting groups. Moreover, 48h after (D2) of the dosing session with ayahuasca, awakening salivary cortisol response (for both sexes) of treated patients became similar to levels detected in controls. This was not observed in patients that ingested placebo that continued showing blunted AUC of awakening salivary cortisol response compared to the control group that ingested placebo. Two days after dosing session (D2), 77% of patients that were treated with ayahuasca showed response while 64% from the placebo group responded. Clinical response was defined as a reduction of 50% or more in baseline scores, of scales of depression.

Cortisol is a steroid hormone that triggers stress response in an adaptive way: it increases cardiovascular and respiratory systems activity, mobilizes glucose to provide enough fuel needed to remove the stressor and limit acute inflammation processes (45). Not only the excess, but also the reduction of this hormone is believed to be harmful, major depression traditionally has been associated mainly with hypercortisolism, but an increasing number of studies have been found hypocortisolism in depression (8, 46, 47). The chronic decrease in cortisol levels induces non-specific symptoms such as general malaise, weakness, low blood pressure, muscle weakness, loss of appetite and weight, gastrointestinal complaints and immunological dysfunction (48). Moreover, low cortisol levels are present in some physiological diseases as Addison’s disease (49) and adrenal insufficiency (50, 51) and some psychopathologies as post-traumatic stress disorders (52). Thus, these results partially corroborate our hypothesis, since control group and patients presented, at baseline, different total plasma cortisol levels and awakening salivary cortisol response, since patients showed hypocortisolemia and blunted awakening salivary cortisol response. But, unexpectedly these alterations in cortisol observed in baseline for patients not were correlated with severity or duration of disease.

In literature more severe depression, marked by a chronic or recurrent disease, frequently disclose hypocortisolism (46, 8, 47). These patients often exhibit long-term exposition to stressors, which in the early phase of disease may induce a chronic upregulated activity of the HPA axis and hypercortisolemia, after this, a maladaptive regulation of the HPA function reduces cortisol to very low pathological levels (53, 54). In addition, evidence suggests that prolonged use of some antidepressants may also lead to increased expression of cortisol receptors and increased sensitivity of negative feedback, thus decreasing cortisol levels to under homeostatic values (55). Our patients are treatment-resistants and used an average of 3.86±1.66 different types of antidepressants, which could in turns induced hypocortisolemia.

Low cortisol levels in depression have also been associated with maladaptive coping style (56) and unfavorable socioeconomic status (57, 13). Patients in this study had a particular profile: they live in Rio Grande do Norte, a state of Northeast Brazil, a region characterized by low socioeconomic status, low educated and living in a low-income household, in a stressful environment. Previous studies, of other research groups, have reported similar results. They found low levels of cortisol at awakening in patients with depression in our state (Rio Grande do Norte) compared to healthy volunteers (58), and with patients from Canada (58). Due to significant levels of poverty and scarce government investments in this region, this population usually is submitted to a long exposure of adverse events in life, and early in life they cope with precarious physical and physiological health, as well as with cumulative social and economic disadvantage conditions.

The etiology of hypocortisolism is explained by various theories. One of them shows that a greater and decompensate sensitivity in negative feedback of the HPA axis deregulates cortisol secretion (54). Also, hypocortisolism is associate to adrenal insufficiency, a failure to produce cortisol and a decrease or increase in Adrenocorticotrophic hormone (ACTH) concentrations that depend on the type of failure, whether primary or secondary, respectively (48). New evidence also appoints the participation of paracrine and autocrine messengers in adrenal failure (59). While several theories try to explain the etiology of hypocortisolism, it seems that the best approach to elucidate this pathophysiological process involves the integration of all these theories.

The regulation of cortisol is an important physiological aspect on the way of achieving biological health, since cortisol is an integrative hormone with large potential of body modulation, particularly involved in the etiology of depression, engaging immune and monoaminergic systems (60, 53). Moreover, optimal levels of cortisol are necessary to induce neurogenesis, possibly due to its modulatory properties over brain-derived neurotrophic factor (BDNF) transcription and binding to its receptor (61, 62, 63). This relation seems an important factor in the etiology of depression, considering that the effectiveness of traditional antidepressants seems to be mediated by neuronal plasticity and neurogenesis (64).

Here, we found that depressive patients showed basal hypocortisolemia and blunted awakening salivary cortisol response in comparison to controls. The values for diagnosis of corticosteroid insufficiency varies, some studies point cortisol levels below 15μg/dl and others under 10μg/dl as hypocortisolemia (65, 66, 67). Studies that evaluated the utility of basal morning serum cortisol measurements in the diagnosis of adrenal insufficiency showed that values below 10μg/dl had 77% of specificity, and 62 of sensitivity (as defined by a subnormal serum cortisol response to insulin-induced hypoglycemia) acting as good indicator of disease (68, 69). In this study we considered 10μg/dl as value of cutoff for true hypocortisolemia. We observed that 61% of patients showed relative hypocortisolemia (below 15μg/dl) and 22.22% true hypocortisolemia (below 10μg/dl). The awakening salivary cortisol response has been less used than plasma cortisol to monitor adrenal insufficiency, because it not had been fully validated as the diagnostic test.

The levels of cortisol in plasma and saliva, at baseline, was positively correlated for controls, but not for patients. It is observed correlation between total plasma and salivary cortisol in healthy subjects (70). Probably this correlation not occurs in patients because of the malfunction of their HPA axis (71) or by changes in concentration of CBG (Cortisol Binding Globulin), its protein of transportation in plasma (70).

During the acute effects of ayahuasca we found increased cortisol levels, 1h40 after intake. Previous studies in healthy subjects have also reported increased cortisol levels during the acute effects of ayahuasca (72, 73, 74), N,N-DMT (75), psilocybin (76,77) and LSD (78). One should bear in mind that our patients presented, in general, hypocortisolemia and blunted awakening salivary cortisol response. It is reasonable to consider that subjects who took ayahuasca which immediately increased cortisol levels probably were acutely benefited by the ingestion of the ayahuasca, thereby leading to a direction of achieving the hormonal homeostasis.

After 48 hours of dosing session, no changes with respect to baseline within each group, both treatments, were observed for AUC of awakening salivary cortisol response and plasma cortisol. Individual changes of AUC and PC between D0 and D2 showed a large variability in these responses. Some studies also faced with this large individual variability in baseline levels and response of cortisol (79), and these are facts that disturb the validation of cortisol as biomarker in DM (80). However, after 48 hours of dosing session the AUC of awakening salivary cortisol response of patients that ingested ayahuasca, and not placebo, became similar to both control that ingested ayahuasca and placebo, the initial blunted response of depressed patients disappear. This similarity of AUC between controls and patients that were treated with ayahuasca points to a beneficial modulation of ayahuasca on awakening salivary cortisol response.

Some studies with animal models of depression, rodents and non-human primates, observed positive antidepressant effects with the use of ayahuasca or its specific components (81, 82, 83). Using the recently validated translational animal model of depression (84), young marmosets (*Callithrix jacchus*) were treated with nortriptyline during 7 days, a tricyclic antidepressant, or with a single dose of ayahuasca. It was observed that ayahuasca increased fecal cortisol levels until 48 hours after it ingestion and presented more notable antidepressant effects than nortriptyline, since it reverted depressive-like behaviors and regulated cortisol levels faster and during more time (83). The modulation of HPA axis by antidepressants depend on the type of antidepressant used and the duration of treatment, acute or chronic. Noradrenaline or serotonin (5HT) reuptake-inhibiting antidepressants, such as reboxetine and citalopram, acutely stimulate cortisol secretion in healthy volunteers, probably due the elevation in 5HT levels (85, 86). On the other hand, some antidepressants, as mirtazapine, acutely inhibits cortisol release, probably due to its selective antagonism at 5-HT2 receptors (85). It is interesting to notice that the long-term effects of antidepressants are frequently opposite of the acute ones. In long way, reboxetine up-regulates cortisol receptors function, repairs the disturbed feedback control and normalizes HPA axis. Mirtazapine, within 1 week, markedly reduces HPA axis activity in depressed patients (85, 86). If the patient is resistant to treatments, and uses antidepressants by years, the long-term effects could be disturbed and followed by a disfavorable physiological response, as cited above, the chronic use of some antidepressant could induces hypocortisolemia.

Here, the acute increases of cortisol levels by the ayahuasca can be due the rise in serotonin induced by the N,N-DMT, and β-carbolinic alkaloids (31, 32, 33), likewise, literature appoints to a modulation of the secretion of CRH and ACTH both at the hypothalamic and pituitary glands by serotonin (87).

In clinical practice the physiological variables are not used for the diagnosis of DM or for choose and evaluate treatments (88). The use of cortisol as a biomarker could aid in the diagnosis, prognosis and analysis of the evolution of the treatments. As discussed, DM is correlated to hyper and hypocortisolemia and antidepressants have distinct action in cortisol levels, thus the use of cortisol as biomarker could influence the choice of treatment. In this way, as ayahuasca increases cortisol levels acutely, its use as antidepressant could be favorable for depressive patients that show hypocortisolemia. However, more investigation are necessary, mainly chronic treatment studies.

Again, once more, our hypothesis was partially corroborated. As was hypothesized, ayahuasca induced a large increase in acute salivary cortisol response than placebo. Although, this increase was not sustained along 48 hours, patients that ingested ayahuasca, and not placebo, presented similar AUC of awakening salivary cortisol response compared to controls (ayahuasca and placebo) in D2. This last finding, although it is different from the hypothesis, is important and corroborates with the improvement of several physiological system, emotional and cognitive aspects that was regulated by cortisol (89, 90). Patients showed considerable reduction in depressive symptoms in D2, however, this progress was not correlated with cortisol changes, either acutely nor 48 hours after dosing.

Our results suggest that the AUC awakening salivary cortisol response is a more robust biomarker than the PC, since it was altered in baseline and it was sensible to treatment by ayahuasca. Other studies appoint in the same way, considering the awakening salivary cortisol response as more strong marker than PC, as it is less modulatated by circadian clock and by daily stressors than PC (79). As well as, some prospective studies have shown that awakening salivary cortisol, and not total PC levels, could predict depressive episodes (91, 92)

However, we should be cautious when trying to consider salivary cortisol, in an isolated way, as a biomarker for the diagnosis of DM, since altered patterns are also found in other mental disorders and in this study we not found correlation between improvement in depressive symptoms and in AUC of awakening salivary cortisol response. Many studies argue that, in an individual way, this biomarker fits more in the aid of prognostics and therapeutic accompaniments than in the diagnosis. On the other hand, it is suggested that a panel of neuroendocrine and immune biomarkers would be the most suitable for the aid in the diagnosis of psychopathologies, as recently proposed for depression (93). In the current overview, however, we emphasize that more studies are required to increase the assumption that salivary cortisol can be useful as a biomarker in order to contribute with valuable information in the diagnostic, prognostic and therapeutic results in major depression

In sum, the present study appoints new evidence of improvements of depressive symptoms and of AUC of awakening salivary cortisol response by ayahuasca, 48 hours after its ingestion, in DM patients with treatment-resistant depression, which presented blunted salivary cortisol awakening response and hypocortisolemia. As cortisol act in regulation of distinct physiological, cognitive and emotional partway, the improvement of its awakening response could be important as part of the antidepressant effects. Taking these findings in account, this work contributes significantly to support the return of clinical studies with natural psychedelics applied to mental disorders.

## Conflict of Interest Statement

That research was conducted in the absence of any commercial or financial relationships that could be construed as a potential conflict of interest.

## Author and Contributors

Galvão-Coelho N., Lobão-Soares B., Araújo D.B. Maia-de-Oliveira, J.P, Arcoverde E and Palhano-Fontes F. designed the experiments; Almeida, R. N. and Galvão A.C measured hormonal data; Freitas, F., Palhano-Fontes F., Onias, H., Galvão A.C. and Silva, E.A. collected experimental data, carried out statistical analysis and prepared figures. Prepared manuscript Galvão-Coelho N., Lobão-Soares B., Araújo D.B., Palhano-Fontes F., Galvão A.C. and Silva, E.A.

## Acknowledgments

The authors would like to express their gratitude to all patients who volunteered for this experiment, and to the Hospital Universitário Onofre Lopes (HUOL), from the Federal University of Rio Grande do Norte, Brazil, for institutional support. This study was funded by the Brazilian National Council for Scientific and Technological Development (CNPq, grants #466760/2014 & #479466/2013), and by the CAPES Foundation within the Ministry of Education (grants #1677/2012 & #1577/2013).

